# Daily light exposure differences between extreme chronotypes persist after alignment to habitual sleep timing

**DOI:** 10.64898/2026.11.11.765960

**Authors:** Johannes Zauner, Manuel Spitschan

**Affiliations:** TUM School of Medicine and Health, Technical University of Munich, Munich, Germany; Max Planck Research Group Translational Sensory & Circadian Neuroscience, Max Planck Institute for Biological Cybernetics, Tübingen, Germany; TUM Institute for Advanced Study (TUM-IAS), Technical University of Munich, Garching, Germany; TUMCREATE Ltd., Singapore, Singapore

**Keywords:** chronotype, melanopic light exposure, photoperiod, circadian entrainment, Munich ChronoType Questionnaire, phase-response curve, dim light melatonin onset

## Abstract

**Purpose:** Morning and evening chronotypes may receive light at different clock times, thereby providing differently timed signals to the circadian pacemaker. Prior studies have suggested phase-specific exposure patterns that could counter further advances in early types and delays in late types. We examined whether photoperiod modifies these exposure patterns in a way that could reasonably counteract timing extremes.

**Methods:** We analysed near-eye melanopic equivalent daylight illuminance (EDI) in 140 participants from an international cohort across seven countries (778 participant-days; 35,949 half-hour observations). We compared four quartiles of midsleep on free days corrected for sleep debt (MSF_sc_) in clock time and relative to MSF_sc_ using cyclic generalized additive mixed models, accounting for day type, site, photoperiod and individual patterns. In exploratory analyses, we tested moderation of these differences by daylength and used phase-response-curve (PRC) weighting as a conceptual comparison of exposure timing.

**Results:** The modelled extreme evening-type/morning-type exposure ratios averaged 0.16 (95% CI 0.06–0.40) 9.6–11.3 hours before MSF_sc_ (pre-MSF_sc_) and 14.0 (5.6–34.8) 0.9–4.4 hours after MSF_sc_ (post-MSF_sc_). The models performed similarly well by ordinary in-sample R² (0.795 for clock time; 0.794 for MSF_sc_-relative time). Pattern differences were broadly similar on work and free days, with the pre-MSF_sc_ contrast descriptively attenuated on free days. The post-MSF_sc_ ratio increased by a factor of 2.59 (95% CI 1.06–6.34) between civil photoperiods one hour below and above each site’s average photoperiod, with no significant moderation of the pre-MSF_sc_ contrast. Exploratory weighting by the published PRC gave higher advance-weighted point estimates in extreme evening than extreme morning types; the group difference was not statistically significant.

**Conclusions:** Extreme chronotypes differ in daily exposure both in clock time and relative to MSF_sc_. The established questionnaire-based MSF_sc_ measure showed significant post-MSF_sc_ daylength moderation. The exposure patterns and conceptual PRC comparison suggest a possible pathway linking daylight availability to entrainment in extreme chronotypes.

## Introduction

Daily timing and light exposure are closely connected: sleep and activity schedules influence when people encounter light, while light contributes to entrainment. Chronotype-related differences in exposure relative to circadian phase are particularly relevant because light can advance or delay the clock depending on when it occurs, as described by a light phase-response curve (PRC).^1–5^ Everyday exposure also depends on the solar day, indoor environments and social schedules.^6^ By comparing groups at the same clock time, we can show when their light environments differ. In turn, comparing them at the same circadian phase asks whether they receive different light input at equivalent points in their biological day.

Previously, Goulet and colleagues compared morning and evening types in clock time and in circadian phase anchored to measured dim-light melatonin onset (DLMO).^7^ Participants were classified into chronotypes using the Morningness–Eveningness Questionnaire (MEQ).^8^ In their sample of 19 participants, the clock-time differences were no longer statistically detectable after DLMO alignment, illustrating how biological time changes the interpretation of light-exposure differences. However, in a subset with extreme DLMO timing, comprising four morning and five evening types, morning types received more light around DLMO, whereas evening types received more light 10–12 hours later. Goulet and colleagues suggested that this exposure pattern could help stabilize entrainment: exposure to light near DLMO could delay early types and counter further advances, whereas light roughly 10–12 hours later could advance late types and counter further delays. This interpretation is consistent with proposed differences in intrinsic circadian period, which the study itself did not measure. Intrinsic period varies between individuals and is associated with morningness–eveningness.^9,10^

Later, Emens and colleagues combined seven days of wrist light measurements with plasma DLMO in 66 MEQ-defined morning and evening types.^11^ They investigated extreme subgroups comprising morning types whose DLMO occurred earlier than every evening type and evening types whose DLMO occurred later than every morning type. They compared light exposure in the advance and delay regions of the PRC. Evening types in these subgroups had a higher advance-to-delay ratio of log-transformed light exposure than morning types, indicating relatively more light in the phase-advancing than phase-delaying region. Their interpretation extended Goulet’s suggestion that phase-specific exposure could help stabilize entrainment.

As an environmental factor, daylight availability might contribute to these differences. When melatonin onset occurs earlier in the solar day, daylight may still be available around that phase, while people with later melatonin onset may reach the same phase after dusk. Natural-light studies show that changes in the light environment can alter circadian timing and that seasonal daylight changes are relevant to human entrainment.^12,13^ Photoperiod is therefore a plausible contributor to exposure that counteracts early or late timing. However, the built environment, electric lighting and behavior shape personal light exposure and can attenuate its seasonal variation.^14,15^ How photoperiod contributes to the previously reported chronotype patterns under everyday conditions remains unclear.

We examined these patterns in our multicentre MeLiDos dataset of personal light exposure across seven countries.^6^ We compared four chronotype groups using local clock time and time relative to midsleep on free days corrected for sleep debt (MSF_sc_), measured using the Munich ChronoType Questionnaire (MCTQ).^16^ MSF_sc_ provides a sleep-based proxy for aligning exposure to circadian phase, distinct from the melatonin-based DLMO measure used by Goulet and colleagues and by Emens and colleagues.^17–19^ We first asked whether chronotype differences remain after this alignment and whether they differ between work and free days. In two post hoc analyses, we then examined how photoperiod modifies the extreme-group contrast and whether weighting by a light PRC places extreme evening types in a more phase-advancing direction than extreme morning types. Together, these analyses extend the entrainment interpretation proposed in the Goulet and Emens studies by testing photoperiod as a possible contributor to the timing of light input.^7,11^

## Methods

### Study and measurements

This analysis used prepared light and diary data from the MeLiDos study and its open-access analysis repository.^6,20^ Measurements were collected from August 2023 to October 2025 at nine sites in seven countries using a common protocol that included ambulatory light measurements, sleep diaries and chronotype questionnaires, among other measures.^21^ Recruitment occurred through the participating local research groups. The preregistration and existing analyses are described elsewhere.^22^ The daylength and PRC analyses reported here were exploratory additions after inspection of the primary chronotype results and were not part of the preregistered analysis.

Participants followed an ambulatory protocol spanning eight calendar days to cover seven days of measurement, with spectrally sensitive ActLumus loggers recording every 10 seconds.^21^ The outcome was melanopic equivalent daylight illuminance (melanopic EDI), which expresses melanopsin-weighted light as the illuminance of an equivalent daylight spectrum, in lux.^23^ Light exposure was measured near the corneal plane using a 3D-printed holder attached to spectacle frames, at the chest using a lanyard and at the wrist. Here, the primary analyses focused on near-eye recordings, and concurrent chest recordings provided a placement sensitivity analysis. During recorded sleep, devices were placed near the bed, so those observations represent the sleeping environment rather than near-eye exposure. Participant-declared waking non-wear was treated as unavailable. For inclusion, we required at least 80% valid coverage of the combined waking and bedside-sleep record per day, excluding wholly zero days. The analysis used prepared 30-minute arithmetic means, requiring at least 15 valid minutes per bin. Remaining zero values were retained, and the response was log10(melanopic EDI + 0.1). This transformation accommodates darkness (0 lux) while retaining variation over the wide range of personal light exposure.^24^

The prepared near-eye data comprised 141 participants and 816 participant-days. Missing MSF_sc_ excluded one participant and six days; missing waking-date diary records excluded a further 32 days. The common sample for both time coordinates comprised 140 participants, 778 participant-days and 35,949 half-hour observations. Participants were 78 female and 62 male, with a median age of 27 years (range 18–68). Work/free-day labels came from the diary for the waking date. [Online Supplement S1](https://tscnlab.github.io/ZaunerSpitschan_JBiolRhythms_2026/analysis.html)^25^ documents source preprocessing, inclusion, model specifications, diagnostics and sensitivity analyses.

### Chronotype and time coordinates

Across the primary dataset, participants were assigned to MSF_sc_ quartiles using one value per person. Boundaries were 03:10, 04:16 and 05:15, and ties were assigned to the earlier quartile. For convenience, we label the quartiles extreme morning, moderate morning, moderate evening and extreme evening, respectively. These labels describe relative positions within this sample. The MEQ was collected alongside the MCTQ, allowing us to assess whether the exposure patterns were consistent across the two chronotype classifications.^8^ Because only three participants were definite evening types, we combined definite and moderate types within each morning-type/evening-type group, excluding intermediate types from this sensitivity only.

We defined clock time as the local midpoint of each half-hour bin. The second coordinate subtracted the participant’s MSF_sc_ and wrapped the result over 24 hours. Negative values therefore precede corrected midsleep. MSF_sc_ was not a daily observation of sleep status: a plotted time after MSF_sc_ does not establish that a participant was asleep or awake then.

DLMO was not measured in this study. For interpretation, we show a DLMO reference interval spanning 6.95–5.95 hours before MSF_sc_. This interval comes from the early- and late-group means reported by Facer-Childs and colleagues in 38 healthy adults (mean age 22.7 years, SD 4.2).^26^ Its midpoint, 6.45 hours before MSF_sc_, anchors our central PRC calculation. The DLMO reference interval does not estimate individual DLMO; questionnaire-based chronotype and DLMO are related but can differ substantially.^18,19^

### Diary midsleep relative to MSF_sc_

In a separate descriptive analysis, we compared MSF_sc_ with the midpoint between diary-reported sleep onset and final awakening. Each sleep episode inherited the work/free-day label of its waking date. We included all valid diary nights from the primary participants, including nights without usable light observations. Pearson correlations were calculated overall and by day type, with 95% intervals from 2,000 participant-level bootstrap samples that retained each person’s nights together. Correlations of participant means provided an equal-participant sensitivity. We also calculated a diary-derived MSF_sc_ equivalent, MSF_sc(diary)_, by correcting mean free-day midsleep for sleep debt using the observed work/free-day night counts:

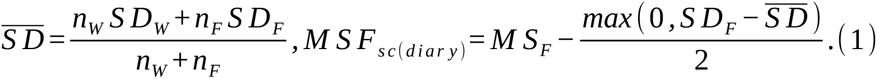

Here, *n_W_* and *n_F_* are the numbers of recorded work- and free-day nights, *S D_W_* and *S D_F_* are their mean sleep durations, and *M S_F_* is mean free-day midsleep. Sleep duration was the interval from reported sleep onset to final awakening. The zero bound in Equation 1 implements the standard conditional correction: sleep debt is corrected only when mean free-day sleep duration exceeds mean work-day sleep duration; otherwise, MSF_sc(diary)_ equals *M S_F_*.^27,28^ Allowing a negative correction would instead shift midsleep later when free-day sleep is shorter, which is not part of this definition.

Our diary adaptation uses recorded mean sleep timing and duration in place of habitual questionnaire estimates, and observed work/free-day counts in place of weekly day counts. Previous studies have also calculated corrected free-day midsleep from sleep diaries.^29,30^ We required at least one night of each type and weighted sleep duration by the observed counts, rather than imposing a five-work-day/two-free-day week. Diary alarm-use information was unavailable, so free nights were identified by the waking day’s work/free label without an additional alarm-use exclusion. Within-person midsleep standard deviations required at least three nights in the relevant day-type context. Calculations used local clock time with midnight handled continuously; [Online Supplement S1](https://tscnlab.github.io/ZaunerSpitschan_JBiolRhythms_2026/analysis.html#sec-sleep-diary)^25^ gives the diary calculations and an elapsed-time sensitivity for daylight-saving transitions.

In a separate sensitivity analysis, we examined whether the findings depended on the chronotype measure and classification. We repeated the exposure models and downstream analyses using MSF_sc(diary)_ and its quartiles. The established questionnaire measure remained primary. A questionnaire-MSF_sc_ repeat on exactly the same eligible participants and light observations separated changes in timing and classification from sample restriction. A further repeat used MSF_sc(diary)_ for phase alignment while retaining the matched questionnaire quartiles, separating the timing change from subsequent reclassification. [Online Supplement S2](https://tscnlab.github.io/ZaunerSpitschan_JBiolRhythms_2026/diary-msfsc-sensitivity.html)^31^ reports these comparisons.

### Daily exposure models and comparisons

We fitted Gaussian generalized additive mixed models separately for clock time and time relative to MSF_sc_. Cyclic smooths represented the common daily pattern, chronotype quartiles, work/free days and their interaction, site differences and individual daily curves. Participant-day intercepts captured day-to-day variation. Common photoperiod terms allowed personal exposure to vary with civil daylength, defined from dawn to dusk at a solar elevation of −6° and centered on each site’s average photoperiod over observed dates. Models were fitted in R 4.6.1 using mgcv and fast restricted maximum likelihood.^32,33^ An AR(1) correction followed actual elapsed time and reset at participant-day boundaries and missing intervals. It also reset around bins whose local clock labels mapped to multiple elapsed-time intervals during daylight-saving transitions; those bins remained in the analysis.

For standardized group predictions, site, participant and participant-day effects were set to zero. Predictions averaged over a common photoperiod distribution, with equal weight per participant and equal weight per observed day within participant. Overall curves used the same pooled work/free-day mixture in every group. Exposure curves were back-transformed standardized log-scale means, not arithmetic mean lux. Their 95% simultaneous bands covered all 12 quartile-by-day-context curves and 144 time points within each coordinate, using the same 10,000 joint coefficient draws and smoothing-parameter-adjusted covariance as the contrast bands below. Personal light exposure patterns were plotted on a symmetric-logarithmic (symlog) axis, linear up to 1 lux and logarithmic above it.^24^

The primary comparisons comprised chronotype direction, chronotype strength, day type and their interactions. We formed 95% simultaneous bands from 10,000 joint coefficient draws over the full family of 20 contrasts and all times within each coordinate, separately from the exposure-curve family. Clock-time comparisons describe differences in the daily light environment; MSF_sc_-relative comparisons ask whether those differences remain after sleep-timing alignment. These separate questions follow the clock-time and phase-aligned comparisons in the literature, so coverage was controlled separately within each coordinate.^7^ We reported intervals only where every contributing quartile/day-type cell contained at least four participants in the local time bin.

Exponentiated log contrasts compare **melanopic EDI + 0.1**. A ratio of one indicates no difference between the compared groups or conditions; a confidence interval excluding one supports a difference. Ratios averaged over an interval are geometric means of the phase-specific ratios. Interval-average 95% confidence intervals used the full coefficient covariance for the mean log contrast, including smoothing-parameter uncertainty, before exponentiation. They treat selected windows as fixed, without multiplicity or selection adjustment, and are distinct from the simultaneous bands used to identify the windows.

We evaluated prediction using five participant-held-out folds and, separately, a held-out final day for eligible participants. Shapley allocations divided the increase in ordinary in-sample R² above the common daily curve among model components by averaging their additional contributions across subset models.^34^ These allocations were conditional on the fitted models and describe their decomposition rather than independent biological contributions. Sensitivity analyses examined chest placement, alternative smooth boundaries, residual-variance weighting and MEQ classification; details are in [Online Supplement S1](https://tscnlab.github.io/ZaunerSpitschan_JBiolRhythms_2026/analysis.html)^25^. Chest and noncyclic comparisons used the same within-coordinate coverage rule. MEQ sensitivity bands jointly covered overall, work-day and free-day contrasts within each coordinate and classification.

### Exploratory daylength moderation

We added a quartile-specific change in the daily curve per hour of site-centered photoperiod to the full MSF_sc_-relative model. This change was linear in photoperiod and common to work and free days. The principal comparison used photoperiods one hour below and above each site’s average photoperiod, with that average as an intermediate reference. We estimated extreme evening-type/extreme morning-type ratios at these targets and their longer-day/shorter-day ratio of ratios. For the latter, a ratio of one denotes no photoperiod moderation; an interval excluding a ratio of one supports a change in the group difference.

Daylength inference used pointwise 95% intervals and interval-average contrasts within the two intervals selected by the primary model. The latter treated those intervals as fixed and were not adjusted for multiplicity or interval selection. Local support required four participants in each extreme-quartile/day-type cell within one hour of the photoperiod target. We also described civil daylight availability relative to MSF_sc_, with equal participant and within-participant day weights. Its shorter/longer absolute-daylength categories were distinct from the within-site-centered model targets. Matched held-out predictions tested whether moderation improved prediction for new participants.

### Exploratory PRC-weighted exposure

To examine light exposure relative to circadian timing, we adapted an ambulatory approach that multiplies daily exposure by a PRC and integrates the result.^35^ We used this weighting as a conceptual comparison of modelled melanopic exposure timing, without predicting phase shifts.

We weighted the primary model’s extreme-group exposure curves by the one-hour bright-white-light PRC of St Hilaire and colleagues.^3^ Numerical coordinates came from the digitized data deposited by Walch and colleagues.^36^ A two-harmonic curve was reconstructed from 18 responses, using pulse midpoint as the time coordinate. We compared the published raw curve, P, with P + 0.54. In the original protocol, phase shifts were assessed from DLMO measured in constant routines separated by three 24-hour cycles with background light below 3 lux. The assumed mean intrinsic period of 24.18 hours implied a free-running delay of 0.54 hours over this interval; adding 0.54 sets that baseline to zero.^3^

Each PRC was divided by its integrated absolute area and used to weight log10(melanopic EDI + 0.1) + 1 across 24 hours. Adding one sets zero exposure to zero on the transformed scale. A positive index indicates that exposure is weighted toward the PRC’s advance region; a negative index indicates weighting toward its delay region. The index is not calibrated in hours of phase shift. The central alignment placed DLMO 6.45 hours before MSF_sc_; the DLMO reference-interval endpoints at 5.95 and 6.95 hours before MSF_sc_ and two wider offsets provided alignment sensitivities. Coefficient-based 95% intervals retained covariance across time and groups, conditional on the reconstructed PRC and assumed alignment. They did not propagate uncertainty in the external PRC or individual circadian phase.

As a sensitivity analysis, we adapted the advance/delay exposure balance of Emens and colleagues.^11^ We applied log10(melanopic EDI + 1) to back-transformed fitted exposure, setting negative EDI predictions to zero. The mean transformed exposure from 16 to 4 hours before DLMO was divided by the mean from 4 hours before to 8 hours after DLMO. We compared balances in extreme evening and morning types by their ratio at the three DLMO reference alignments, using 10,000 joint coefficient draws for unadjusted 95% percentile intervals conditional on the fitted model and assumed DLMO.

## Results

### Sample and chronotype distribution

Our four MSF_sc_ groups contained 36, 34, 36 and 34 participants, respectively (Figure 1). The extreme groups contributed 70 participants and 390 participant-days. MEQ scores overlapped across MSF_sc_ quartiles, illustrating that sleep timing and morning/evening preference do not give identical classifications. The MEQ sensitivity analysis retained 68 participants and 379 participant-days after excluding 72 intermediate participants.

**Figure 1:**
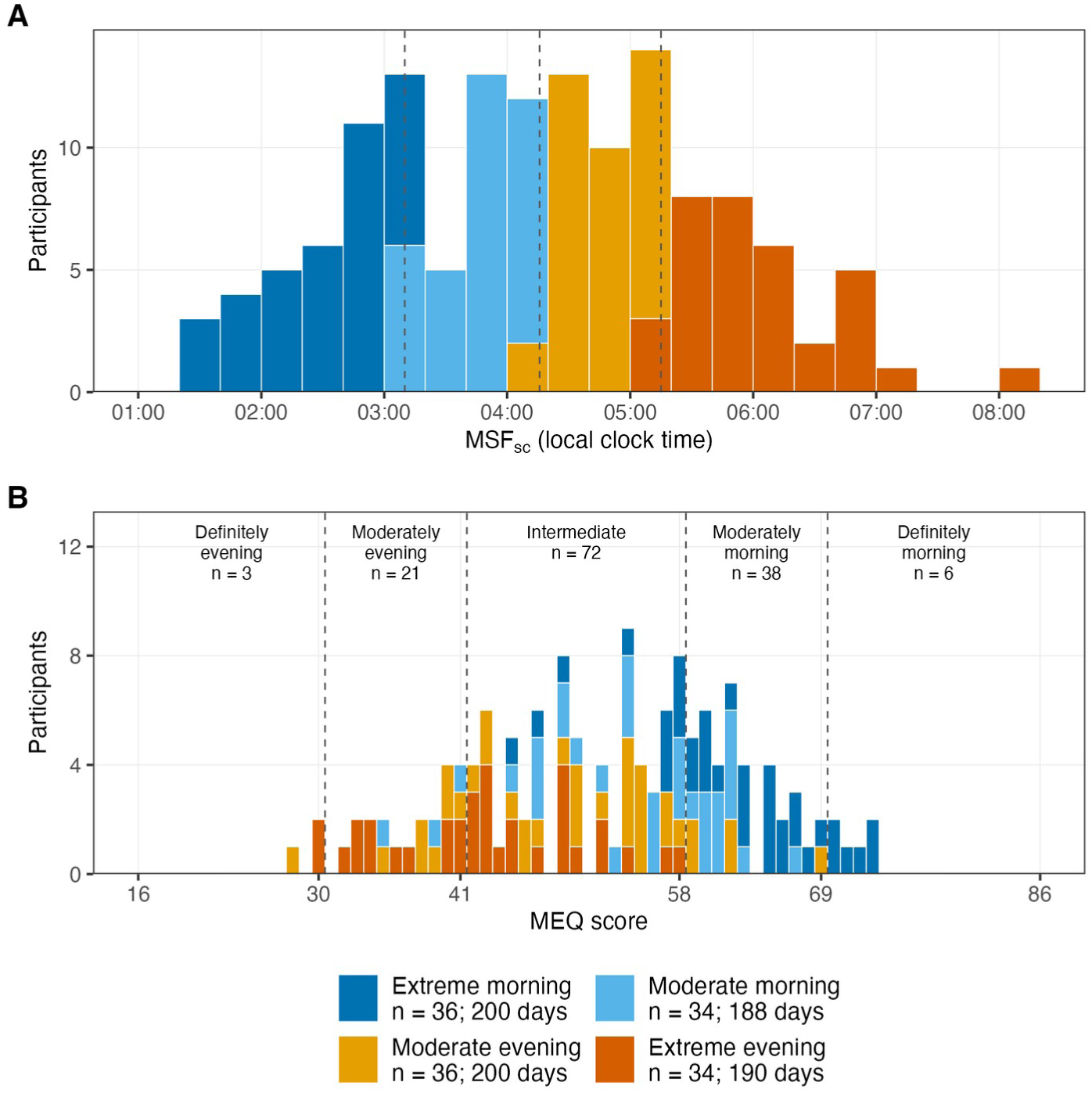
Chronotype distributions among 140 participants. (A) MSF_sc_ in 20-minute bins, with quartile boundaries at 03:10, 04:16 and 05:15. (B) MEQ scores in one-point bins, with category boundaries. Colors indicate the same MSF_sc_ quartiles in both panels. Each participant contributes once; participant-day counts describe available exposure records and do not weight the histograms.

### Diary midsleep corresponds with MSF_sc_ but varies across nights

The 140 primary participants contributed 971 valid diary nights: 620 before work days and 351 before free days (Figure 2 A). Diary midsleep correlated with MSF_sc_ overall (r = 0.66; 95% CI 0.59–0.73), on work days (r = 0.67; 0.59–0.74) and on free days (r = 0.74; 0.67–0.81). Correlations of participant means were 0.81, 0.75 and 0.82, respectively (Figure 2 B). The stronger correlations after averaging nights fit MSF_sc_’s role as a measure of habitual sleep timing rather than an individual night’s midpoint. The stronger free-day association is consistent with MSF_sc_ being based on free-day midsleep, with work-day sleep duration contributing to its sleep-debt correction.

**Figure 2:**
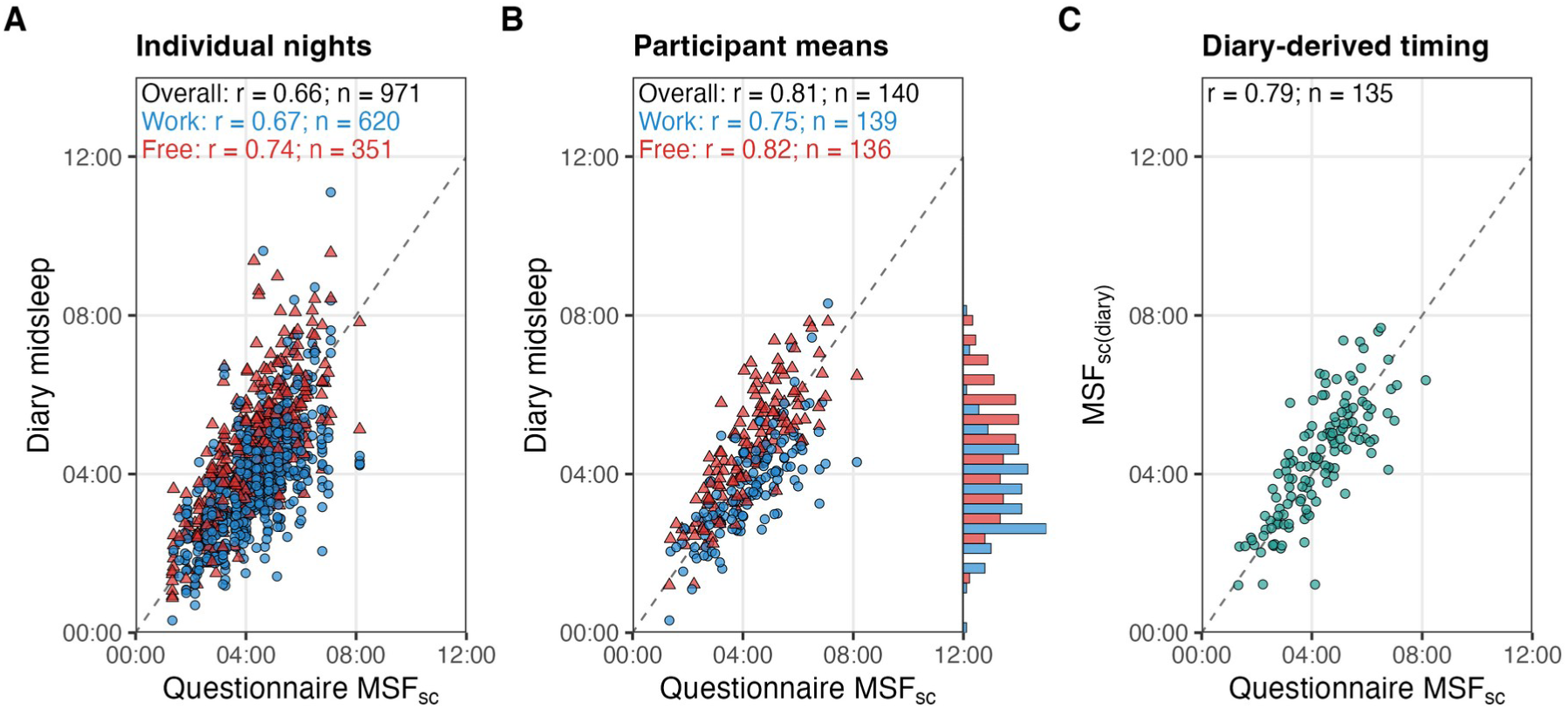
Diary sleep timing and questionnaire MSF_sc_. (A) Individual nights from 140 participants: 620 before work days and 351 before free days. (B) Participant-average midsleep, with 139 work-day and 136 free-day means. Blue circles and labels denote work days; red triangles and labels denote free days. The adjacent histogram counts these means in common 30-minute bins; the two day types occupy offset half-width bars within each bin. Overall correlations in A and B use all nights and one all-night mean per participant, respectively. (C) Diary-derived corrected midsleep (MSF_sc(diary)_) for 135 participants with both day types, using observed work/free-day counts in the sleep-debt correction. All times are local clock times; dashed diagonals mark equality. Labels give Pearson correlations and n: nights in A, participants in B and C.

Among the 139 participants with at least three nights available, the median within-person midsleep SD was 47 minutes (IQR 34–62). Diary midsleep was typically earlier than MSF_sc_ before work days and later before free days, with median differences of −34 and +18 minutes. At the group level, these modest offsets follow the expected work/free-day directions and do not suggest a large, uniform displacement from MSF_sc_.

For the 135 participants with both work-day and free-day nights, MSF_sc(diary)_ correlated with questionnaire MSF_sc_ at r = 0.79 (95% CI 0.72–0.85; Figure 2 C). Its median difference was +6 minutes (IQR −28 to +36), compared with +23 minutes for uncorrected mean free-day midsleep in the same participants. The correction reduced the typical offset, although the correlation was slightly lower than for the uncorrected free-day means (r = 0.82). [Online Supplement S1](https://tscnlab.github.io/ZaunerSpitschan_JBiolRhythms_2026/analysis.html#sec-sleep-diary)^25^ gives day-type variability and results using all available participants.

### Alignment to MSF_sc_ broadly reverses the clock-time differences

In clock time, exposure increased and declined earlier in morning types than in evening types (Figure 3 A). Alignment to MSF_sc_ broadly reversed the extreme-group differences in these portions of the daily pattern (Figure 3 B). Extreme morning types had higher fitted exposure approximately 9.6–11.3 hours before MSF_sc_ (pre-MSF_sc_), with an average extreme evening-type/extreme morning-type ratio of 0.16 (95% CI 0.06–0.40). Extreme evening types had higher exposure approximately 0.9–4.4 hours after MSF_sc_ (post-MSF_sc_), with an average ratio of 14.0 (95% CI 5.6–34.8). Ratios during the rising portion of the curve can be large even when absolute exposure is low.

**Figure 3:**
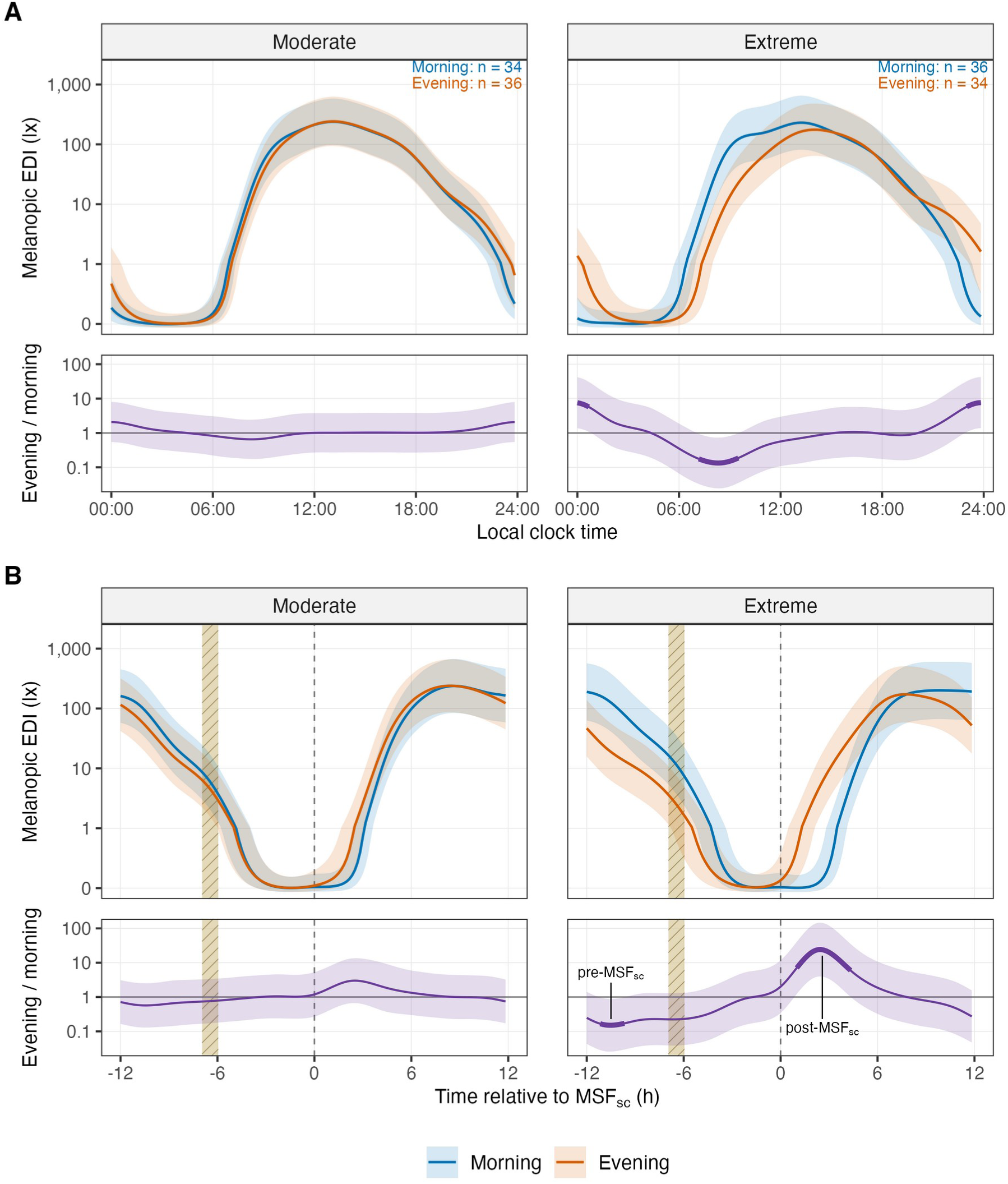
Daily melanopic light exposure and evening-type/morning-type contrasts. (A) Local clock time. (B) Time relative to MSF_sc_. Columns compare moderate and extreme morning-type and evening-type groups. In each block, upper panels show back-transformed standardized means of log10(melanopic EDI + 0.1), with 95% simultaneous bands; shorter lower panels show evening-type/morning-type ratios of melanopic EDI + 0.1. Curves use a common photoperiod distribution and work/free-day mixture, excluding site and individual effects. The exposure axis is linear up to 1 lux and logarithmic above it. A ratio of one indicates equal exposure between the compared groups; a band excluding one supports a difference. Within each time coordinate, the upper bands have 95% simultaneous coverage over all 12 quartile-by-day-context curves and 144 time points; the lower bands separately cover all 20 registered contrasts over the same time points at 95%. Bold lavender segments exclude one and meet the primary local-support rule of four participants per contributing quartile/day-type cell. Centered black labels in B identify the significant extreme-group intervals, with pre-MSF_sc_ above and post-MSF_sc_ below their intervals; short vertical black leaders touch neither labels nor curves. Dashed vertical lines mark MSF_sc_ at 0 h in the exposure and ratio panels of B; the striped band is the DLMO reference interval.

Pattern differences were broadly similar on work and free days. The pre-MSF_sc_ evening-type/morning-type ratio was 0.13 on work days and 0.21 on free days, indicating descriptive attenuation on free days. The corresponding post-MSF_sc_ ratios were 15.5 and 12.0. Simultaneous interaction bands did not show a significant difference in the chronotype contrast between day types.

The MEQ sensitivity analysis reproduced the post-MSF_sc_ pattern, with higher evening-type exposure. For the overall comparison, alternative noncyclic group curves retained a significant post-MSF_sc_ difference, whereas the pre-MSF_sc_ band included one. The post-MSF_sc_ contrast was therefore more stable than the pre-MSF_sc_ contrast to the spline specification ([Online Supplement S1](https://tscnlab.github.io/ZaunerSpitschan_JBiolRhythms_2026/analysis.html#sec-meq)^25^).

### Fit and prediction

Clock-time and MSF_sc_-relative models performed similarly well by ordinary in-sample R², 0.795 and 0.794, respectively. These fit statistics include participant and day effects. Chronotype direction and strength together received Shapley allocations of 0.007 R² in the clock-time model and 0.012 R² in the MSF_sc_-relative model, above the common daily curve. Individual daily patterns received the largest allocation. Alignment to MSF_sc_ therefore exposed interpretable group differences without making chronotype the dominant source of variation.

Using clock time gave slightly better predictions for new participants: predictive R² was 0.668, compared with 0.652 for time relative to MSF_sc_, and log-scale root mean squared error was 0.887 versus 0.908. The paired root mean squared error difference was 0.0205 (95% CI 0.0108–0.0310). Prediction of a held-out final day for known participants was similar, with predictive R² of 0.674 for clock time and 0.672 for time relative to MSF_sc_.

### Daylength moderation is clearest post-MSF_sc_

The extreme-group ratio post-MSF_sc_ increased with photoperiod (Figure 4 A–C). Across the primary post-MSF_sc_ interval, the geometric mean evening-type/morning-type ratio was 7.99 at the −1-hour target and 20.69 at the +1-hour target. Their ratio was 2.59 (95% CI 1.06–6.34). In the pre-MSF_sc_ interval, the ratio of ratios was 0.83 (0.33–2.09), providing no clear evidence of moderation. A separate trough around 8.33 hours before MSF_sc_ had a pointwise estimate of 0.41 (0.16–1.04). This interval included a ratio of one, indicating no significant change in the group difference with photoperiod at this phase.

**Figure 4:**
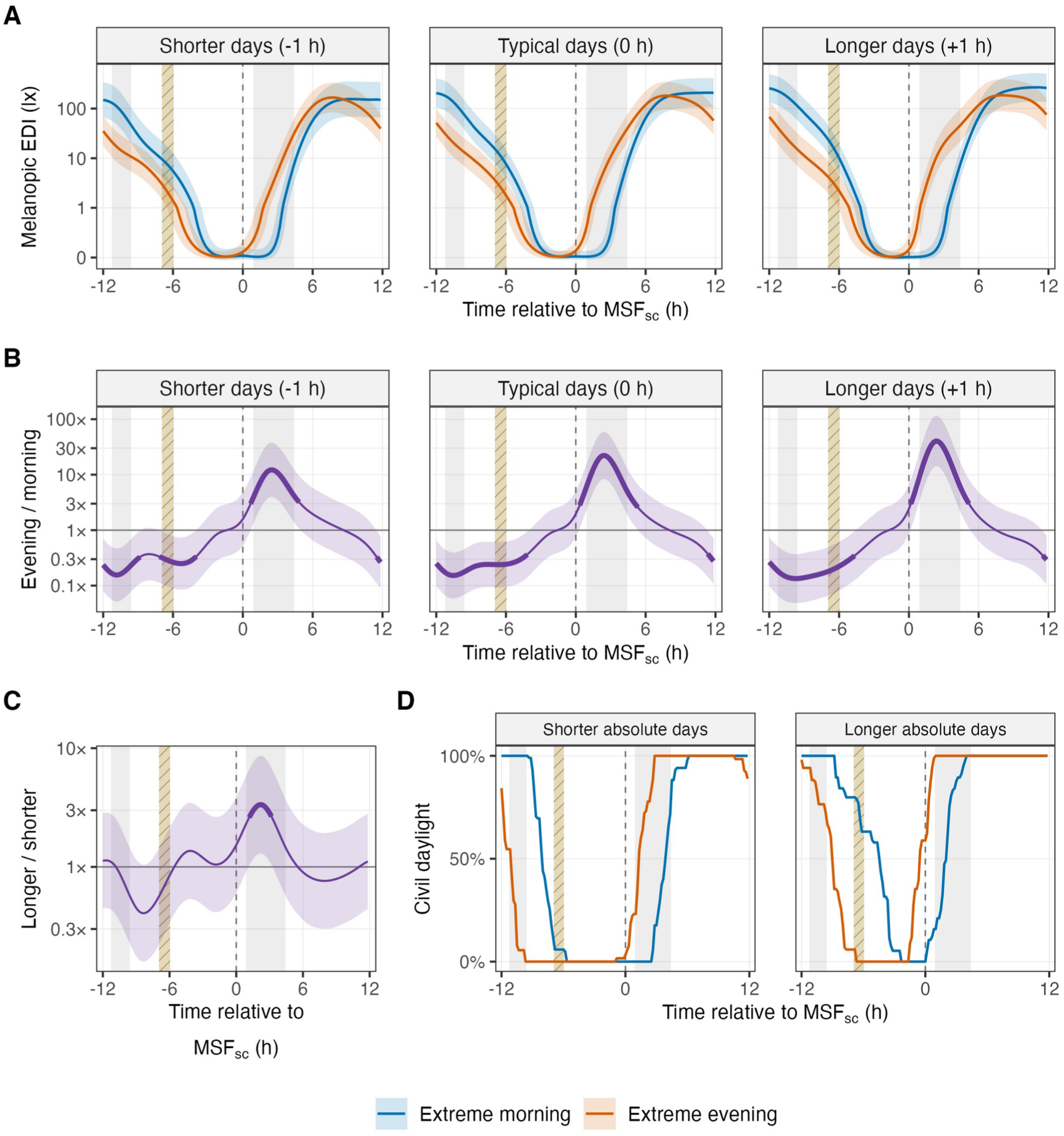
Exploratory daylength moderation and astronomical daylight availability. (A) Extreme-group curves at photoperiods one hour below, at, and one hour above each site’s average photoperiod over observed dates. (B) Corresponding extreme evening-type/extreme morning-type ratios of melanopic EDI + 0.1. (C) Longer-day/shorter-day ratio of those group ratios. A ratio of one indicates no change in the evening-type/morning-type contrast between photoperiod targets; a band excluding one supports moderation. Bands in A–C are pointwise 95% intervals. Bold segments in B–C exclude one and meet the exploratory local-support rule of four participants per contributing extreme-quartile/day-type cell. (D) Civil daylight availability relative to MSF_sc_ among 70 extreme-group participants contributing 390 days, with equal participant and within-participant day weighting. The shorter/longer absolute-daylength categories in D differ from the within-site-centered targets in A–C. Dashed vertical lines mark MSF_sc_ at 0 h in all panels. Grey shading without boundary lines marks the two primary intervals; the striped band shows the DLMO reference interval. Civil photoperiod runs from dawn to dusk at a solar elevation of −6°.

In the extreme groups, astronomical daylight occurred at different MSF_sc_-relative times (Figure 4 D). Within the pre-MSF_sc_ interval, civil daylight was more available to morning types; within the post-MSF_sc_ interval, it was more available to evening types. These availability gaps were smaller on longer absolute days. These descriptive differences in exposure opportunity do not directly translate to the within-site-centered moderation of measured exposure (Figure 4 A–C).

Adding photoperiod moderation did not improve new-participant prediction. Predictive R² was 0.649 for the expanded model and 0.652 for the primary MSF_sc_-relative model. The paired root mean squared error difference was 0.0044 (95% CI −0.0008 to 0.0100).

### Evening types trend toward a more phase-advancing exposure pattern than morning types

Our PRC-weighting analysis showed that, at the central DLMO alignment, overall indices based on the published raw PRC were −0.152 (95% CI −0.351 to 0.046) for extreme morning types and 0.046 (−0.148 to 0.240) for extreme evening types (Figure 5). Both intervals spanned zero. With the drift-corrected PRC, indices were 1.276 (1.104–1.448) and 1.422 (1.255–1.589), respectively, indicating advance-weighted exposure in both groups.

**Figure 5:**
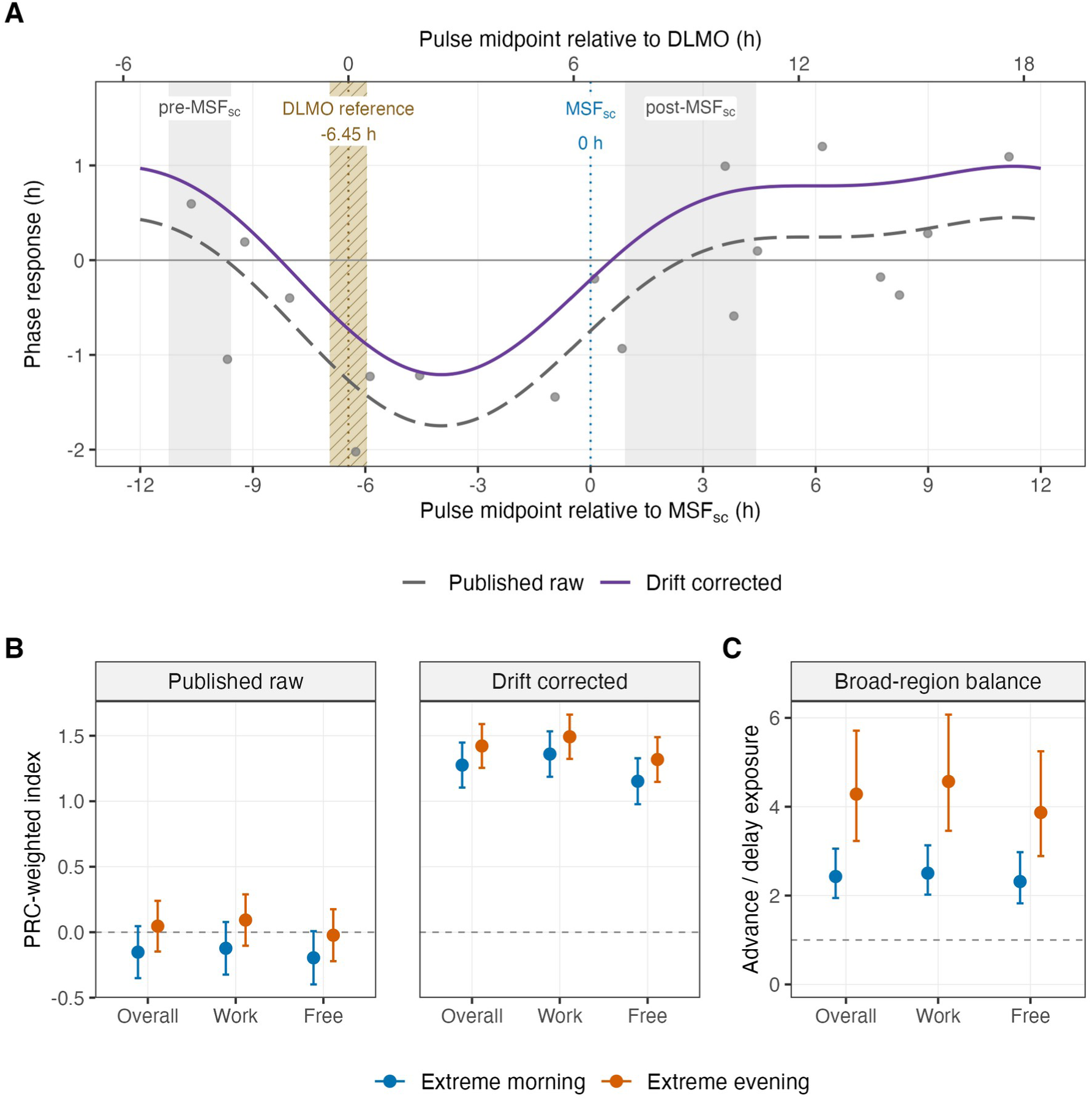
Exploratory PRC weighting and advance/delay sensitivity of extreme-group exposure. (A) Two-harmonic reconstruction of the one-hour bright-white-light PRC of St Hilaire et al.,^3^ using coordinates digitized and deposited by Walch et al.^36^ Points are 18 responses from the PRC study. Pulse midpoint is aligned to DLMO at −6.45 h relative to MSF_sc_. The dashed curve preserves the published raw baseline; the solid curve adds 0.54 h for drift correction. Curves precede absolute-area normalization. (B) Weighted indices under the raw and drift-corrected PRCs for extreme morning-type and evening-type groups at the central alignment, overall and separately for work and free days. Exposure is zero-exposure-centered as log10(melanopic EDI + 0.1) + 1. Bars show 95% coefficient intervals conditional on the PRC and alignment. Positive values indicate advance-weighted exposure balance, not phase shifts in hours. (C) Emens-style advance/delay balances using log10(fitted melanopic EDI + 1) at the central alignment, overall and by day type, with 95% percentile intervals. One denotes equal regional means within a group; one for the evening-type/morning-type ratio reported in the text would denote equal balances between groups. Extreme groups comprise 70 participants and 390 participant-days. In A, labelled grey bands identify the pre-MSF_sc_ and post-MSF_sc_ intervals, and the striped band shows the DLMO reference interval.

The extreme evening-type minus morning-type contrast was positive under both baselines: 0.198 (95% CI −0.094 to 0.491) with the raw PRC and 0.146 (−0.106 to 0.398) with drift correction. Every direct group-contrast interval included zero at the central alignment, overall and separately on work and free days. Across the DLMO reference alignments at 5.95, 6.45 and 6.95 hours before MSF_sc_, only the raw-PRC work-day contrast at 6.95 hours before MSF_sc_ narrowly excluded zero, 0.297 (0.002–0.593); all overall contrast intervals included zero. The point estimates consistently indicated more advance-weighted exposure in evening types than morning types, or more delay-weighted exposure in morning types than evening types.

At the central DLMO alignment, the Emens-style advance/delay balances were 2.43 (95% CI 1.94–3.06) in extreme morning types and 4.28 (3.23–5.71) in extreme evening types (Figure 5 C). The evening-type/morning-type ratio of balances was 1.76 (95% CI 1.21–2.58); its overall interval remained above one at both DLMO reference interval endpoints.

### Diary-derived MSF_sc_ preserves exposure differences but weakens daylength moderation

To test whether the findings depended on questionnaire-based sleep timing, we repeated the analyses using diary-derived MSF_sc(diary)_ for phase alignment and recalculated the chronotype quartiles.

The matched questionnaire and MSF_sc(diary)_ analyses included 135 participants, 755 participant-days and 34,884 half-hours; 50 participants changed quartile. The most common transitions were between the two evening-type groups: from extreme to moderate (11 participants) and from moderate to extreme (9 participants); [Online Supplement S2](https://tscnlab.github.io/ZaunerSpitschan_JBiolRhythms_2026/analysis.html#sec-sleep-diary)^31^ gives the full classification matrix. Within the original phase windows, the extreme evening-type/morning-type ratio was 0.13 (95% CI 0.05–0.33) pre-MSF_sc(diary)_ and 28.3 (11.3–70.9) post-MSF_sc(diary)_, compared with 0.18 (0.07–0.46) and 14.7 (5.8–37.7) using questionnaire MSF_sc_ on the same observations. These unadjusted interval-average comparisons retained the direction of both exposure differences.

The post-MSF_sc_ longer-day/shorter-day ratio was 2.66 (95% CI 1.10–6.43) in the matched questionnaire analysis. Using diary timing while retaining questionnaire quartiles gave 2.39 (0.95–6.02); also assigning quartiles from MSF_sc(diary)_ reduced the estimate to 1.15 (0.52–2.54). Thus alignment alone left the point estimate close to the questionnaire result, although its interval included one. The larger reduction in the point estimate followed reclassification. Both exposure contrasts retained their directions with fixed questionnaire groups: 0.29 (0.11–0.76) pre-MSF_sc_ and 8.7 (3.5–21.7) post-MSF_sc_.

The relative PRC-weighted evening-type direction persisted in the full MSF_sc(diary)_ analysis: at the transferred central DLMO alignment, the raw-PRC contrast was 0.306 (0.008–0.604), whereas the drift-corrected contrast was 0.236 (−0.021 to 0.494). [Online Supplement S2](https://tscnlab.github.io/ZaunerSpitschan_JBiolRhythms_2026/diary-msfsc-sensitivity.html)^31^ reports alignment, prediction, attribution and model-structure comparisons.

## Discussion

We detected significant exposure differences between extreme morning and evening types in both clock time and MSF_sc_-relative time, but no significant differences between moderate morning and evening types. Aligning exposure to MSF_sc_ broadly reversed the extreme-group differences seen in clock time. Morning types had higher exposure early in the local day, whereas evening types had higher post-MSF_sc_ exposure. Morning-type exposure also declined earlier in clock time, yet was higher in the pre-MSF_sc_ interval. The post-MSF_sc_ difference was broadly similar on work and free days and remained visible under an alternative chronotype questionnaire. Exploratory daylength modeling also linked the average post-MSF_sc_ contrast to photoperiod. Together, these results support considering daylight availability alongside individual sleep timing when describing chronotype-specific exposure.

The diary comparison provides convergent support for using MSF_sc_ as a reference for habitual sleep timing. Participant-average midsleep correlated more strongly with MSF_sc_ than individual nights, and the modest work/free-day offsets followed the expected directions. Applying the sleep-debt correction to the diary means also brought their typical timing closer to questionnaire MSF_sc_, with a median difference of six minutes. This correspondence is consistent with Suh and colleagues’ seven-day diary comparison, which found an association between diary- and questionnaire-derived MSF_sc_ in young adult good sleepers.^29^ The median within-person SD of 47 minutes nevertheless shows that actual midsleep varies around this reference. Consequently, the pre- and post-MSF_sc_ exposure patterns describe group differences relative to habitual corrected midsleep while averaging over variation in nightly sleep schedules. The post-MSF_sc_ interval broadly corresponds to the later part of habitual sleep, but does not identify the second half of every recorded sleep episode. Near awakening, this variation can change whether a given MSF_sc_-relative exposure occurs during sleep or wakefulness.

Diary-derived MSF_sc_ depends on the observation schedule and the definition of free days. Sasawaki and colleagues required five school nights and two non-school nights with awakening without an alarm.^28^ Winnebeck and colleagues used diary averages with fixed five-school-day/two-free-day weighting.^30^ Our observed-count weighting describes the recorded period, which may differ from the habitual week if coverage is uneven. A single free night can also give an unstable estimate, and a work-free day does not guarantee unconstrained awakening when alarm use is unknown. These differences support our treatment of MSF_sc(diary)_ as a complementary, period-specific measure rather than an interchangeable replacement for questionnaire MSF_sc_. Despite the lower minimum-night requirement, diary-derived and questionnaire estimates were strongly correlated (r = 0.79).

The established questionnaire-based MSF_sc_ measure is the basis of the primary analysis. The alternative approach based on sleep-diary-derived MSF_sc(diary)_ retained both directions of the exposure contrast but did not show significant daylength moderation. The same-participant questionnaire analysis retained significant moderation, so sample restriction alone does not explain this difference. Holding questionnaire quartiles fixed while changing alignment to MSF_sc(diary)_ largely preserved the estimated moderation, although its interval included one. Reclassifying chronotypes produced the larger reduction in the point estimate in this sequential comparison. The exposure-pattern directions were more stable across these specifications than the daylength association.

Later chronotypes encounter the dawn transition earlier relative to MSF_sc_. The civil-daylight summaries show more daylight for extreme morning types pre-MSF_sc_ and extreme evening types post-MSF_sc_, matching the direction of both measured exposure differences. This timing agreement supports a daylight contribution to the post-MSF_sc_ pattern. The pre-MSF_sc_ interval-average comparison showed no significant photoperiod moderation. Gaps in astronomical daylight availability between morning and evening types narrowed on longer absolute days, whereas the average fitted evening-type/morning-type ratio increased with site-centered photoperiod. Indoor attenuation, artificial light and behavior determine how astronomical opportunity becomes measured exposure.

Our findings complement natural-light camping studies, in which later chronotypes showed larger phase advances after a week outdoors, and seasonal and weekend changes in the light-dark environment altered melatonin timing.^12,13^ Those interventions changed the light-dark environment within participants; our observational data compare everyday exposure between participants observed at different photoperiods. The camping interventions did not isolate a post-MSF_sc_ photoperiod effect. Further, a mathematical model predicts that brighter daytime light narrows the distribution of circadian phase and sleep timing, with a stronger effect at longer photoperiods.^37^ These findings provide context for the post-MSF_sc_ association observed here.

Comparing our findings with those of Goulet and colleagues is informative, but does not constitute a direct replication.^7^ Their extreme-DLMO subset showed higher evening-type exposure at a later biological phase, qualitatively consistent with the post-MSF_sc_ result here. Their morning-type excess near DLMO was not matched by a significant extreme-group interval within our DLMO reference interval; our pre-MSF_sc_ difference lay before it. Our larger sample and near-eye melanopic measurements extend the evidence from Goulet and colleagues’ wrist light recordings, while their measured DLMO provided a more direct metric of circadian phase than our sleep-based proxy.

Emens and colleagues interpreted the greater advance-to-delay exposure balance in evening types as potentially meeting the greater daily advance needed to entrain a longer intrinsic circadian period.^11^ This interpretation concerns the light input required to maintain stable entrainment, rather than evidence of progressive phase advancement. Our Emens-style sensitivity supported the relative evening-type/morning-type direction using broad, equally weighted phase windows.

Previously, an ambulatory pilot study used a related approach, applying a nonlinear intensity–response transformation to personal illuminance before multiplying it by the Khalsa PRC and integrating advance and delay areas.^35^ Our use of log-transformed melanopic exposure, the one-hour PRC and MSF_sc_-based alignment is an adaptation of that approach. This precedent supports the weighting operation as a conceptual summary of exposure timing; it does not calibrate our index as a phase-shift prediction.

Our PRC weighting places extreme evening-type exposure in a relatively more phase-advancing direction and extreme morning-type exposure in a relatively more phase-delaying direction, but the direct group-contrast intervals included zero at most DLMO reference alignments. In this weighting approach, the raw-PRC group intervals spanned zero, while drift correction yielded positive indices for both groups. The index also depends on exposure level and the reference assigned a neutral, zero contribution on the centered log scale (here 0 lux), because each PRC has nonzero signed area. Laboratory PRCs describe responses to particular stimuli, including one-hour white-light, 6.7-hour white-light and 6.5-hour short-wavelength exposures.^2–4^ Phase resetting also varies nonlinearly with exposure duration.^38^ Summing weighted exposure therefore does not establish that naturally varying light acts additively. Dynamic circadian models and regressions using light in advance/delay windows have predicted measured DLMO from ambulatory records, but these calibrated models differ from a fixed PRC-weighted integral.^39^ Validating our index as a phase-shift predictor would require comparison with measured changes in circadian phase under corresponding exposure patterns. Here, it remains a conceptual summary of exposure timing.

The relatively small allocation of fitted variation to chronotype and the absence of a predictive gain from daylength moderation put the group findings in context. Individual patterns accounted for much more of the fitted variation. Aligning to MSF_sc_ is useful for examining when groups differ relative to their own sleep timing, while clock time retained slightly better prediction for new people. This is compatible with the group results: a chronotype contrast can identify a consistent difference during part of the day while accounting for little of the total variation that determines overall prediction error.

This analysis has several limitations. The observational design cannot determine whether light exposure shaped chronotype, chronotype shaped exposure, or both reflected schedules and environments. A week of observation per participant also cannot show how an individual’s exposure pattern changes across seasons.

Finally, near-eye recordings approximate the light environment without measuring retinal exposure, and bedside recordings during sleep do not incorporate eyelid transmission. DLMO and circadian phase shifts were not measured. A stronger test of the daylight explanation would observe both extreme groups at the same sites across seasons and measure DLMO. Such a design could, together with a separation of outdoor, indoor and sleep-associated exposure, distinguish daylight opportunity from behavior and test whether the post-MSF_sc_ contrast translates into different circadian responses.

We asked whether chronotype differences persist after MSF_sc_ alignment, vary with photoperiod and favor different phase-shifting directions. Extreme morning types had higher pre-MSF_sc_ exposure, whereas extreme evening types had higher post-MSF_sc_ exposure. With the established questionnaire-based MSF_sc_ measure, photoperiod was significantly associated with the post-MSF_sc_ contrast, with no significant pre-MSF_sc_ moderation. The alternative approach based on sleep-diary-derived MSF_sc(diary)_ did not show significant daylength moderation. The main PRC analysis suggested a relatively more phase-advancing evening-type pattern with uncertain group contrasts; our Emens-style sensitivity supported that relative direction. Together, these findings are compatible with a role for light exposure and photoperiod in moderating further advances or delays in extreme chronotypes, helping maintain stable 24-hour entrainment.

## Supplementary material

[Online Supplement S1](https://tscnlab.github.io/ZaunerSpitschan_JBiolRhythms_2026/analysis.html)^25^ contains the complete computational workflow, sample registries, model diagnostics, work/free-day and placement comparisons, MEQ and smooth-basis sensitivities, prediction and attribution analyses, daylength and PRC sensitivity results, and the diary-midsleep comparison. [Online Supplement S2](https://tscnlab.github.io/ZaunerSpitschan_JBiolRhythms_2026/diary-msfsc-sensitivity.html)^31^ repeats the calculations using diary-derived timing alongside a matched questionnaire analysis and the fixed-quartile comparison. The supplements, source snapshots, package versions and exported results are available in the [chronotype analysis repository](https://github.com/tscnlab/ZaunerSpitschan_JBiolRhythms_2026), with released snapshots preserved in the [Zenodo archive](https://zenodo.org/records/24143023).^40^

## Acknowledgements

We thank the MeLiDos participants and the collaborating site teams for making these recordings available.

## Statements and Declarations

### Ethical considerations

The source study was approved by the Ethics Committee of the Technical University of Munich for Tübingen (2023-115-S-KK) and Munich and Dortmund (2024-118-S-SB); the Swedish Ethical Review Authority for Borås (2024-06792-01); the THUAS Ethics Committee for Delft (positive review, 29 May 2024); the Research Ethics Committee of Universidad San Pablo-CEU for Madrid (834/24/106); the Scientific Research and Publication Ethics Committee of the Faculty of Science and Engineering at İzmir Institute of Technology (decision 5/1); the Comité Ético Científico of Universidad de Costa Rica for San José (CEC-279-2025); and the Committee on Human Research, Publications and Ethics of Kwame Nkrumah University of Science and Technology for Kumasi (CHRPE/AP/644/24).^6^

### Consent to participate

All participants provided written informed consent.

### Academic, standardisation and professional roles

Member of Joint Technical Committee 20 (JTC20) of the International Commission on Illumination (CIE); member of the Research Data Alliance Working Group on Optical Radiation Exposure and Visual Experience Data; member of DIN working committee FNL 27 (NA 058-00-27 AA), “Effect of light on human beings”, within the DIN Standards Committee Lighting Technology; member of the specialist group on melanopic light effects (FG 02) within the Technical-Scientific Committee (TWA) of the German Society for Lighting Technology and Lighting Design (LiTG), and former spokesperson of FG 02; and member of LiTG’s Expert Forum for Interior Lighting (EFI).

### Remunerated roles

Examiner for the Swiss Lighting Society; teacher for LiTG, Munich University of Applied Sciences, and Rosenheim Technical University of Applied Sciences; software developer for LiTG; associated partner at 3lpi lighting design + engineering, Munich; tool and 3D-model designer for Zumtobel Lighting GmbH; and course designer for Munich University of Applied Sciences and Virtual University Bavaria.

### Honoraria for talks

LiTG; Lamilux/Heinrich Strunz GmbH; Robert-Bosch Hospital Stuttgart; Ergotopia GmbH; the German statutory accident insurance institution for the administrative sector (VBG); BRIXEN CULTUR; KITEO GmbH & Co. KG; and the University of Applied Sciences Augsburg.

### Travel reimbursements

Daimler und Benz Stiftung and BG ETEM.

### Intellectual property

Together with 3lpi, holder of a design patent for a non-visually optimized luminaire, No. 008194021–0001 through −0006, at the European Union Intellectual Property Office.

Manuel Spitschan declares the following potential competing interests between 2021 and 2025: academic roles as a member of the Board of Directors of the Society of Light, Rhythms, and Circadian Health, Chair of Joint Technical Committee 20 of the International Commission on Illumination, member of the Daylight Academy, and Chair of the Research Data Alliance Working Group Optical Radiation and Visual Experience Data; remunerated roles as Speaker of the Steering Committee of the Daylight Academy, ad hoc reviewer for the Health and Digital Executive Agency of the European Commission and the Swedish Research Council, Associate Editor for *LEUKOS*, examiner for the University of Manchester, Flinders University and the University of Southern Norway, and consultant for LyS Technologies and RoX Health; research funding and support from the Max Planck Society, Max Planck Foundation, Max Planck Innovation, Technical University of Munich, Wellcome Trust, National Research Foundation Singapore, European Partnership on Metrology, VELUX Foundation, Bayerisch-Tschechische Hochschulagentur, BayFrance/Bayerisch-Französisches Hochschulzentrum, BayFOR/Bayerische Forschungsallianz and Reality Labs Research; honoraria for talks from ISGlobal, the Research Foundation of the City University of New York and Stadt Ebersberg, Museum Wald und Umwelt; travel reimbursements from the Daimler und Benz Stiftung; and being named on European Patent Application EP23159999.4A, “System and method for corneal-plane physiologically-relevant light logging with an application to personalized light interventions related to health and well-being.” Manuel Spitschan declares that these roles and relationships had no influence on the work presented here.

## Data availability

The source MeLiDos datasets are available through the [MeLiDos Project GitHub organisation](https://github.com/MeLiDosProject) and archived in the [MeLiDos Zenodo community](https://zenodo.org/communities/22nrm05_melidos/records?q=&l=list&p=1&s=10&sort=newest). The [source analysis repository](https://github.com/tscnlab/ZaunerEtAl_bioRxiv_2026) provides the original analysis code and prepared data. The accompanying chronotype analysis contains a fixed input snapshot, source checksums, R code and package versions for reproducing this analysis, available in the [chronotype analysis repository](https://github.com/tscnlab/ZaunerSpitschan_JBiolRhythms_2026). Released repository snapshots are preserved in the [Zenodo archive](https://zenodo.org/records/24143023), doi:10.5281/zenodo.24140946.40

## Artificial-intelligence assistance

OpenAI Codex Desktop version 26.623.141536 (build 4753), using GPT-6 Astra (gpt-6-astra, Max reasoning setting), assisted with R and Quarto code development, reference checks, numerical checks, figure and table preparation, and manuscript drafting and revision. The authors take responsibility for all calculations, interpretations, and text.

